# RNA cytometry of single-cells using semi-permeable microcapsules

**DOI:** 10.1101/2022.09.24.509327

**Authors:** Greta Leonaviciene, Linas Mazutis

**Affiliations:** Institute of Biotechnology, Life Sciences Centre, Vilnius University, 7 Sauletekio av., Vilnius, LT-10257, Lithuania

**Keywords:** semi-permeable microcapsules, droplet microfluidics, single-cell RT-PCR, multiplex PCR, RNA cytometry, coacervates

## Abstract

Analytical tools for gene expression profiling of individual cells are critical for studying complex biological systems. However, the techniques enabling rapid measurements of gene expression on thousands of single-cells are lacking. Here, we report a high-throughput RNA cytometry for digital profiling of single-cells isolated in liquid droplets enveloped by a thin semi-permeable membrane (microcapsules). Due to selective permeability of the membrane, the desirable enzymes and reagents can be loaded, or replaced, in the microcapsule at any given step by simply changing the reaction buffer in which the microcapsules are dispersed. Therefore, complex molecular biology workflows can be readily adapted to conduct nucleic acid analysis on encapsulated mammalian cells, bacteria, or other biological species. The microcapsules support sequential multi-step enzymatic reactions, and remain intact under different biochemical conditions, freezing, thawing and thermocycling. Combining microcapsules with conventional FACS provides a high-throughput approach for conducting RNA cytometry of individual cells based on their digital gene expression signature.

## INTRODUCTION

The reverse-transcription polymerase chain reaction (RT-PCR) remains one of the most commonly used molecular biology techniques for a quick evaluation of gene expression levels in cells and tissues ^1,2^. The extraordinary sensitivity of one nucleic acid molecule per 1 μl reaction volume ^3^, combined with high specificity, accuracy and fast readout offered by RT-PCR assays have propelled a broad range of biological and diagnostic applications ^2,4-7^. However, a comprehensive characterization of heterogeneous biological samples ^8-10^, such as tumors, microorganisms, or viruses, requires isolation of each individual member in a population so as to study it separately. As a result, numerous efforts have focused on developing single-cell RT-PCR (scRT-PCR) approaches using microtiter plates ^11-14^, microfluidics ^15-28^ and other reaction formats ^29-31^. For example, conventional fluorescence-activated cell sorting (FACS) into microtiter plates followed by RT-PCR ^12,32-35^, enables the analysis of gene expression of sorted cells relative in to their surface markers. While clearly useful for some biological and clinical applications ^34,35^, unfortunately, such approach has not found broader use due to the limited number of single-cells that can be simultaneously processed in 96-well plates and the relatively high costs associated with large reaction volumes ^36^. Moreover, the unbiased characterization of complex populations may require the profiling of thousands of cells individually at a scale that is beyond the practical scope of plate-based assays ^37-39^. FISH-flow enables a high-throughput single-cell profiling, yet sample preparation process is time-consuming (>30h) and is prone to cells loss and RNA degradation ^40^. Alternative techniques based on microwells and valve-operated microfluidics ^41,42^ offer the throughput of scRT-PCR assays of up to 1,000 reactions per run ^43^, but such assay formats often rely on sophisticated microfluidic operations ^21^ that are difficult to implement, prone to errors and expensive ^44,45^.

Droplet microfluidics technology provides an alternative for performing ultra-high-throughput assays at a scale of 10^6^ reactions per run ^41,46^. Individual cells can be isolated in pico-to nano-liter volume range droplets, lysed, and their genetic makeup can be assessed using various molecular biology techniques ^24,25,37^. Therefore, combining the high sensitivity of the RT-PCR technique with a scalability of droplet microfluidics technology, opens a possibility of developing digital assays for high-throughput quantification and analysis of the transcriptional profile of single-cells at a molecular level. However, a digital scRT-PCR assay that is broadly applicable in biological and biomedical research must fulfill several criteria. First, it should provide an efficient approach to isolate and retain encapsulated cells throughout all analytical procedures. Second, the method should enable complex biochemical reactions on thousands of single-cells simultaneously, including efficient cell lysis, nucleic acid amplification and analysis in a straightforward manner using a regular laboratory pipette. Third, the digital scRT-PCR signal should efficiently differentiate true positive events (cells), from false positive events (e.g., cell-free nucleic acids), and other confounding outcomes. Finally, although not critical, the compartments should provide an option to release the amplified genetic material for further analysis, without damaging. To the best of our knowledge, no high-throughput, single-cell RT-PCR technique reported to date fulfills the above requirements, although several noticeable examples that rely on complex microfluidic operations have been reported ^17,19,25^.

Here, we introduce a novel high-throughput RNA cytometry approach for digital profiling of single-cells isolated in liquid droplets with a semi-permeable membrane (microcapsules). The concept is easy to appreciate: the mammalian cells are isolated in liquid droplets whereby each droplet, on average, contains one cell. The liquid droplets, comprising dextran and chemically-modified oligopeptide, form a liquid core (enriched in dextran) and a liquid shell (enriched in oligopeptide), the later which is cross-linked into a thin and thermostable membrane. Once microcapsules are formed, the subsequent analytical procedures are carried out using a regular pipette and tubes (i.e., no need for expensive equipment). We showed that using microcapsules as reaction vessels enables efficient isolation and retention of mammalian cells, and sustains the multi-step analytical procedures required for the extraction, purification, amplification and digital analysis of nucleic acids. The microcapsules readily withstand various chemical environments (e.g., solvents), freezing and thermocycling, and are compatible with conventional FACS used to conduct high-throughput, single-cell RNA cytometry, and to accurately quantify the gene expression of thousands of individual mammalian cells.

## RESULTS

### Microcapsules for high-throughput nucleic acid analysis of individual cells

The overall concept of microcapsule-based approach for conducting single-cell RNA cytometry is summarized in Figure 1. At first, the mixture of cells is isolated in biocompatible microcapsules having a liquid core surrounded by a thin semi-permeable membrane. The encapsulated cells are then lysed by dispersing the microcapsules in a lysis mix and processed through a series of washing steps to purify the cells’ genetic material. The size-selective permeability of a membrane prevents cellular nucleic acids from escaping the compartments, while simultaneously enabling the intracellular proteins (e.g., RNases) and other low-molecular-weight compounds to leave the microcapsule. Therefore, RNA and DNA molecules encoded by individual cells can be efficiently purified and retained within the compartments. Once the nucleic acids are purified, the microcapsules are transferred to the RT-PCR reaction mix to initiate cDNA synthesis, followed by multiplex PCR. During PCR, the fluorescently labelled primers in the reaction mix cross the membrane by diffusion and are incorporated into the PCR amplicons, rendering the compartments with a cell fluorescent. Thus, using a multiplex panel of fluorescently labelled PCR primers, the expression of the selected genes of interest — in hundreds to tens of thousands of individual cells — can be digitally profiled and quantified using a conventional flow cytometry, or regular epifluorescence microscope.

**Figure 1.**
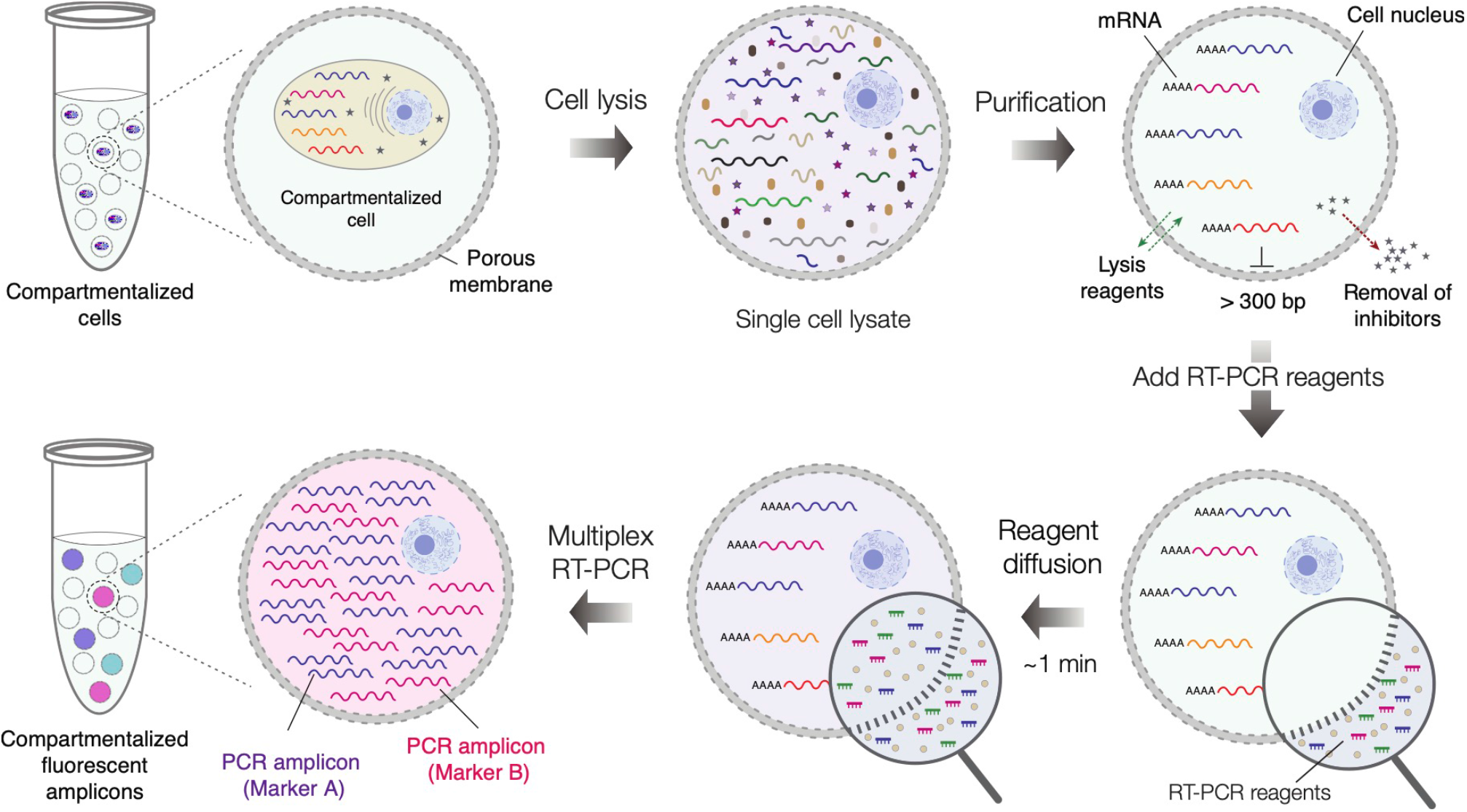
The concept of performing single-cell multiplex RT-PCR using microcapsules. The individual cells are isolated (compartmentalized) in semi-permeable microcapsules and are lysed by dispersing microcapsules in an appropriate lysis buffer. Upon lysis the nucleic acid molecules longer than 300 bp. are retained within the microcapsules, while low molecular weight biomolecules are removed by dialysis. Following cell lysis step, the microcapsules carrying purified nucleic acids are dispersed in RT-PCR reaction mix containing fluorescently labelled oligonucleotides. Once dispersed in a reaction mix, the reaction components quickly diffuse to the microcapsule enabling multiplex RT-PCR reaction on encapsulated nucleic acids. After RT-PCR the microcapsules carrying a particular cell type, and the transcript(s) of interest, become fluorescent thus enabling differentiation of encapsulated cells based on their digital transcriptional profile.

### Generation of uniform microcapsules for efficient mammalian cell isolation and retention

To realize the experimental strategy presented in Figure 1, we generated microcapsules with a semi-permeable membrane composed of a gelatin derivative, a thermo-responsive oligopeptide that solidifies at lower temperatures and can be crosslinked into a 3D gel mesh. To produce uniform microcapsules, we first generated aqueous two-phase system (ATPS) droplets in the carrier oil on a 40-μm deep co-flow microfluidics device at a throughput of 540 droplets s^-1^, by infusing gelatin methacrylate (GMA) and dextran solutions, with the cells suspended in the latter at a desirable density (Figure 2A). Following droplet generation at room temperature, the GMA and dextran phases quickly phase-separated, forming a liquid shell enriched in GMA and a liquid core enriched in dextran (Figure 2B). The shell surrounding the liquid core was converted to an elastic membrane in a two-step procedure. First, the shell was solidified by cooling the droplets to 4 ºC and then covalently cross-linked under a brief exposure to a light-activated photo-initiator (see *Materials and Methods*). This two-step polymerization procedure, where a physical gelation of microcapsules is followed by a covalent cross-linking, ensured a highly reproducible generation of microcapsules comprising a well-centered core, enriched in a liquid dextran, and an elastic membrane composed of covalently cross-linked oligopeptide. We confirmed that microcapsules remain intact under variety of experimental conditions such as freezing, thawing, centrifugation at high speeds (e.g., 20,000 rcf), and in the presence of different salts and solvents.

**Figure 2.**
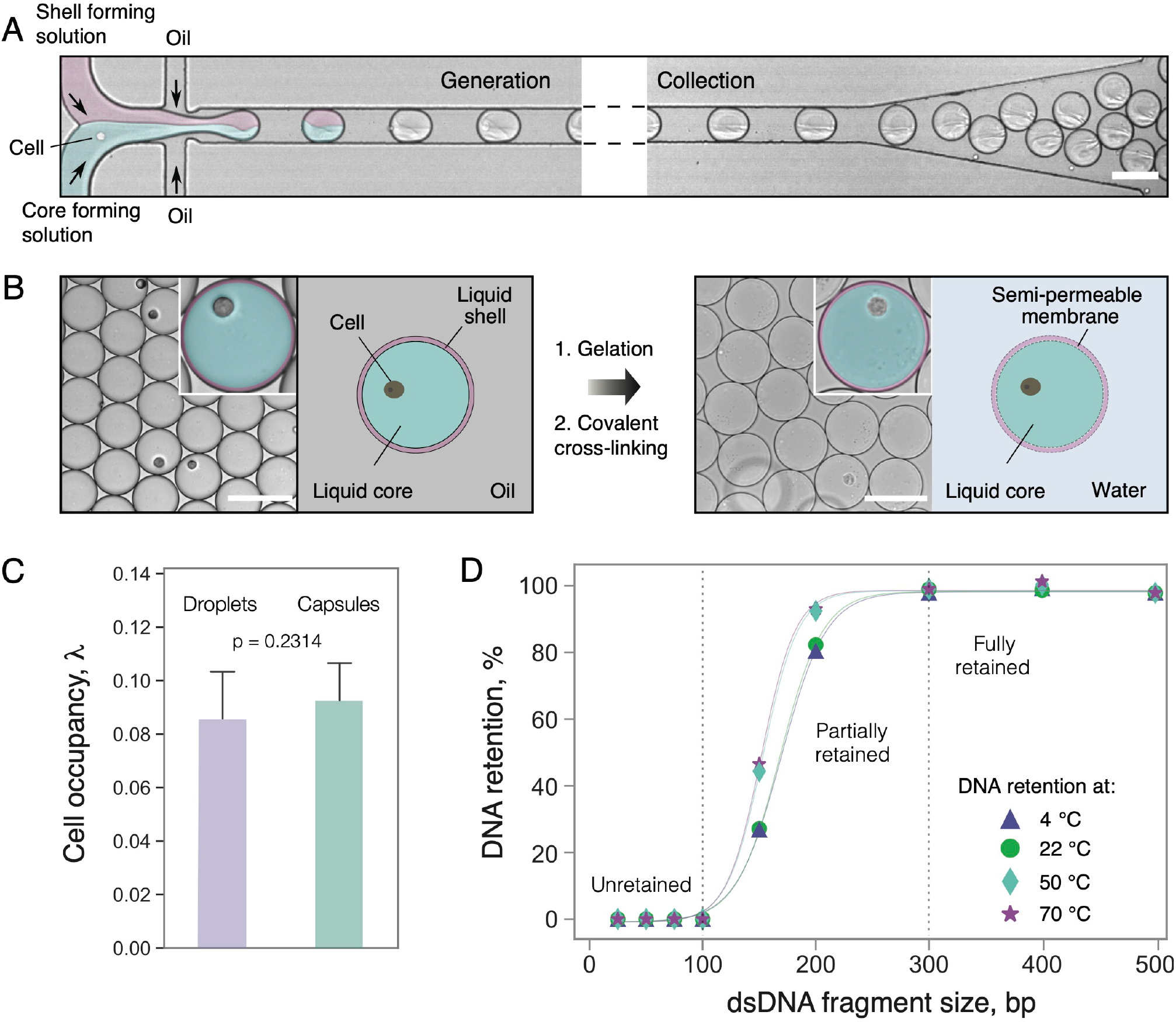
Microcapsule generation, single-cell encapsulation and DNA retention. A) Cell encapsulation and collection off-chip. A mixture of cells is encapsulated in aqueous two-phase system (ATPS) droplets comprising a liquid core and a liquid shell and collected off-chip in the tube. B) The liquid shell of collected droplets is converted to a semi-permeable membrane in a two-step process: at first, the liquid shell is gelled by cooling, and then covalently cross-linked by photo-polymerization. C) The boxplots representing mammalian cell retention in droplets and microcapsules. D) DNA retention in microcapsules at different temperatures after 30 min incubation. The DNA fragments 100 bp. and smaller freely transverse the microcapsule’s membrane, while DNA fragments 300 bp. and longer are fully retained. Solid lines serve only for a visual guidance. Scale bars, 100 μm.

To evaluate the mammalian cell encapsulation and retention efficiency, we generated 75.0 ± 1.5 μm size microcapsules with a 3.0 ± 0.2 μm thick membrane. We loaded human cells (K-562) and quantified cell retention microscopically immediately after encapsulation, and after forming the microcapsules. No significant difference was detected (two-tailored *t*-test (n = 18) *t* = 1.22, *p* = 0.2314) confirming that compartmentalized cells are efficiently retained during the microcapsule generation (Figure 2C). These results sharply contrast with hydrogel bead–based assays, which experience uncontrolled cell loss due to the cell’s tendency to adhere at the hydrogel-oil interface ^47,48^. Hence, the use of microcapsules, composed of dextran and GMA, ensures minimal or no cell loss.

### Nucleic acid retention and permeability of microcapsules

While efficient cell isolation and retention are necessary for conducting single-cell assays, to be broadly applicable in RT-PCR the microcapsules must also ensure efficient retention of nucleic acid molecules during the analytical procedures. To experimentally verify the permeability and retention of the nucleic acid molecules within microcapsules, we encapsulated dsDNA fragments ranging from 25 to 700 bp. and incubated the microcapsules at temperatures ranging from 4 to 70 ºC for 30 minutes. We then washed the microcapsules in a neutral pH buffer to remove all unretained DNA molecules and extracted the retained-DNA within the microcapsules by dissolving the membrane (see *Materials and Methods*). The results, shown in Figure 2D clearly indicate that DNA fragments ≥300 bp. were efficiently retained inside the microcapsules, the molecular weight of which approximately corresponds to ∽180 kDa. Considering the typical oligonucleotide length of 20– 50 nt. (12–30 kDa), and the molecular weight of M-MLV reverse transcriptase (71 kDa) and Taq DNA polymerase (94 kDa), we postulated that microcapsules with such permeability are well-suited for conducting massively parallel biochemical reactions such as RT-PCR on thousands of single-cells by simply dispersing them in a suitable reaction mix.

### Digital multiplex RT-PCR enables accurate cell type identification

To illustrate the digital profiling of individual human cells based on their gene expression, we mixed human leukemia cell line (K-562) and human embryonic kidney cells (HEK293) at a 1:1 ratio, and encapsulated the mixture at a dilution level such that each microcapsule, on average, would contain no more than one cell. As a reference, we also separately encapsulated either K-562 or HEK293 cells. We first lysed the encapsulated cells by dispersing the microcapsules in a chaotropic lysis mix followed by extensive washes and genomic DNA (gDNA) digestion with DNase I to obtain the microcapsules with purified total RNA derived from single-cells. The microcapsules were then dispersed in the RT reaction mix to convert the cellular mRNA to copy DNA (cDNA) using poly(dT) primers (see *Materials and Methods* for further details). To identify individual cells based on their digital expression signature, the post-RT microcapsules were transferred into a PCR reaction mix supplemented with fluorescently labelled primers targeting cell-specific markers (Supplementary Table S1). The cDNA of transcripts that encode the protein tyrosine phosphatase receptor type C (PTPRC) and yes-associated protein 1 (YAP) were chosen as targets of interest for the K-562 and HEK293 cells, respectively, whereas the cDNA of β-actin (ACTB) served as a reference. The target-specific PCR primers were fluorescently labelled at 5’-end with fluorophores emitting light at different wavelengths, to enable identification of amplified nucleic acids based solely on the fluorescence signal. During the PCR, the fluorescently labeled oligonucleotides diffused from the bulk solution into the interior of the microcapsules and upon binding to the target DNA template got incorporated into the PCR amplicon thereby transforming an amplified DNA into a fluorescent product (Figure 3). Only the microcapsules carrying a target template (e.g., PTPRC, YAP or ACTB) could incorporate the fluorescent probes into newly synthesized amplicons rendering them fluorescent (Figure 3B). The microcapsules lacking nucleic acid template remained blank. As a result, the microcapsules containing a cell of interest could be distinguished by a cell-type-specific fluorescence signal. For example, given the differential expression of PTPCR and YAP genes ^49^ and the ubiquitous expression of ACTB, the post-RT-PCR microcapsules harboring K-562 and HEK293 cells turned positive in two channels, PTPCR-ACTB and YAP-ACTB, respectively (Figure 3C, cyan and magenta). Altogether these results demonstrate that microcapsule-based scRT-PCR assay enables accurate cell-type identification based on their digital gene expression signature.

**Figure 3.**
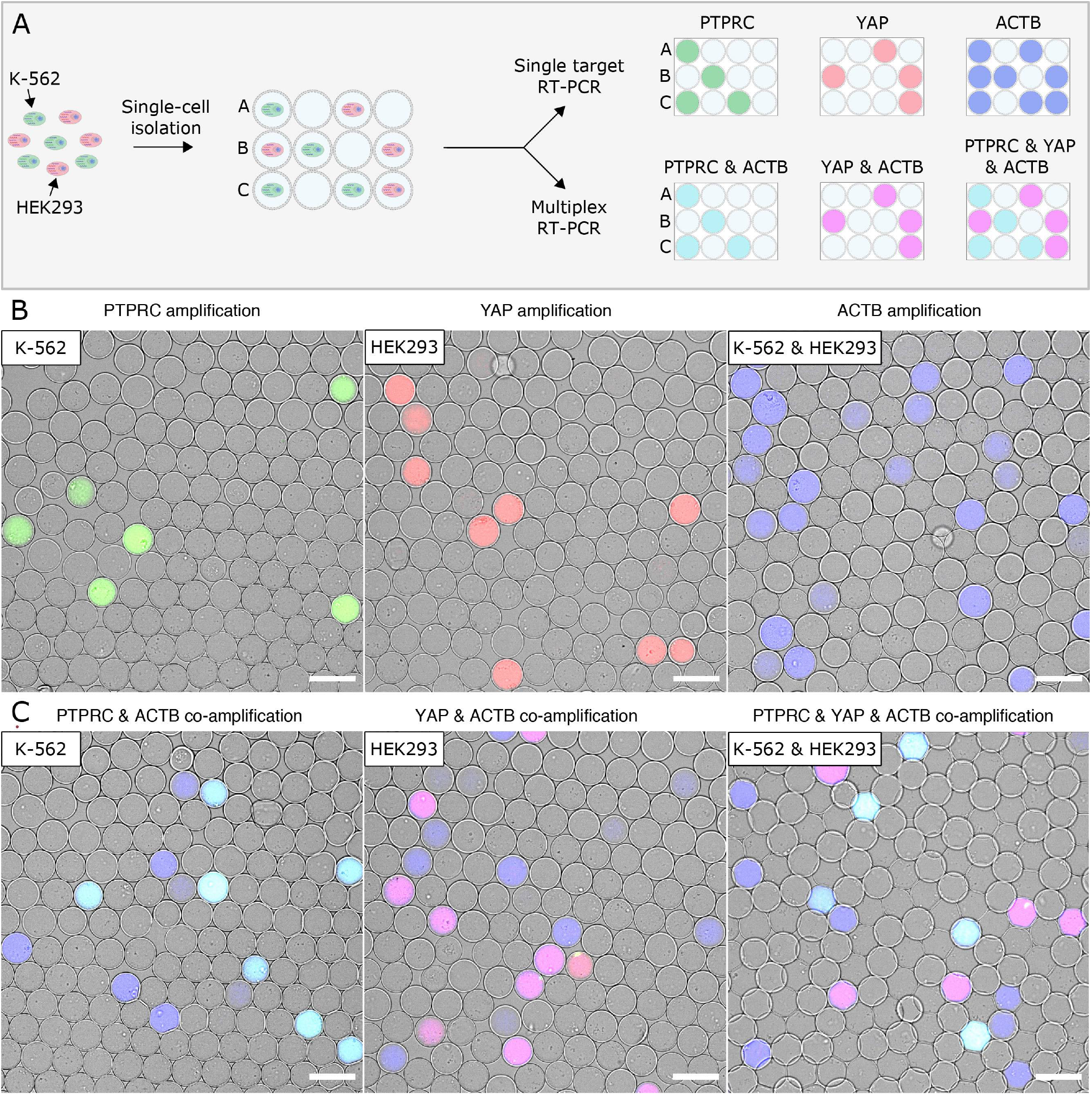
Digital gene expression profiling using microcapsule-based RT-PCR. A) Schematic of the experimental design. The mixture of K-562 and HEK293 cells is loaded in microcapsules and subjected to uniplex RT-PCR or multiplex RT-PCR assay using fluorescently labelled primers targeting the transcripts of interest (PTPRC, YAP, ACTB). The digital gene expression profile of individual cells is then analyzed microscopically by layereing the microcapsules on a hemocytometer and recording their fluorescence. B) Epifluorescence images of microcapsules after RT-PCR reaction targeting one transcript of interest: PTPRC (green), YAP (red) or ACTB (blue). C) Epifluorescence images of microcapsules after multiplex RT-PCR reaction simulteniously targeting two or three transcripts of interest: PTPRC and ACTB (cyan), YAP and ACTB (magenta) or PTPRC, YAP and ACTB (cyan/magenta). Scale bars, 100 μm.

### Flow cytometry for a high-throughput analysis of intrinsic cell markers

To validate the microscopy results presented in Figure 3 and eliminate potential under-sampling artifacts, we explored the high-throughput capabilities offered by flow cytometry instruments. We loaded the post-RT-PCR microcapsules carrying K-562 cells, HEK293 cells, or a mixture of both, onto the FACS instrument and performed RNA cytometry (Figure 4). As a demonstration, up to 30,000 microcapsules were analyzed per experiment, although the total microcapsule count was not limited and could be easily scaled up. The distribution of flow cytometry events in the forward vs. side scatter plot (Figure 4, Capsule gate), fluorescence of Alexa Fluor 647 vs. side scatter plot (Figure 4, Actin gate) and Alexa Fluor 488 vs. Alexa Fluor 555 intensity plot (Figure 4, Cell marker gate) enabled precise quantification of cell-type-specific marker expression and identification of different cell types in the population. The results showed a very close agreement between the flow cytometry and epifluorescence microscopy measurements. The microcapsules carrying K-562 cells were PTPRC positive (3.19–3.63% events), whereas the microcapsules carrying HEK293 cells were YAP positive (4.84–5.46% events), approaching the theoretical frequency of cell dilution (∽5-6%). The microcapsules prepared with a mixture of HEK293 and K-562 cells showed a positive signal either in the PTPRC or YAP channel, while K-562 and HEK293 co-encapsulation events were rare (0.02–0.15%), and followed the Poisson distribution. Measuring co-expression of one for marker gene and one for ubiquitously expressed gene, facilitated the digital profiling of single-cells. For instance, 99.07% of PTPRC-positive events were also positive for the ACTB signal in the sample containing K-562 cells, and 94.57% of YAP-positive events were also positive for the ACTB signal in the sample containing HEK293 cells. Hence, the microcapsules are fully compatible with FACS instruments and enable a high-throughput cytometry of cells based on their gene activity rather than surface proteins based on antibody staining.

**Figure 4.**
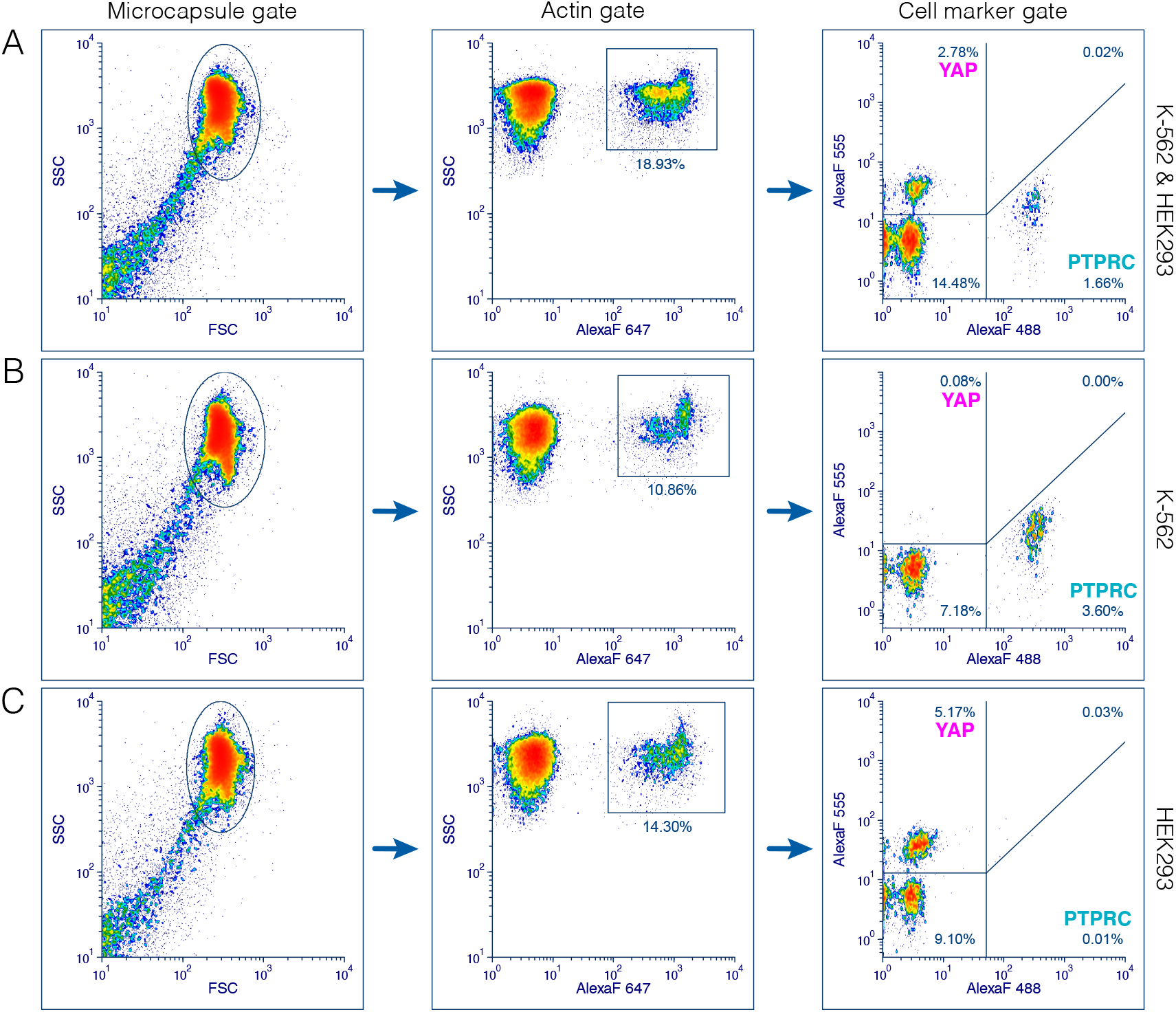
Flow cytometry of individual cells based on intrinsic cell-specific marker expression. The FACS plots show three microcapsule samples containing a mixture of K-562 and HEK293 cells (A), K-562 cells (B) or HEK293 cells (C). The microcapsules were first gated based on forward vs. side scatter signal (Microcapsule gate), the resulting sub-population was then gated on ACTB marker expression (Actin gate) and finally the expression of PTPRC and YAP markers was evaluated on the fluorescence scatter plot (Cell marker gene). The percentage indicates the total events.

Intriguing, the comparative flow cytometry and microscopy analysis revealed the presence of a third subpopulation with a fluorescent signal corresponding to the ACTB target alone and lacking PTPCR and YAP expression. A fraction of ACTB-positive events was approximately 3-times higher than the number of target-specific YAP or PTPRC events. Considering the large excess of ACTB-positive events as compared to the number of loaded cells, we postulated that these events may represent ambient RNA molecules that were present in the initial cell suspension due to premature cell lysis. This notion is supported by the previous reports showing that scRT-PCR in water-in-oil droplets leads to an increased fraction of false positives than the actual number of cells ^17,25^, and that the ambient RNA released during cell preparation is the most likely source of these events ^27^. To rule out the possibility that ACTB-positive events are a by-product of nucleic acid molecule exchange during RT-PCR (e.g., RNA or cDNA molecules escape the microcapsule containing a cell and then enter neighboring empty microcapsule where it gets amplified) we performed a control experiment in which 2 μg of purified total RNA from K-562:HEK293 cell mixture was added to the RT-PCR mixture supplemented with ∼200.000 blank (empty) microcapsules. In case there is nucleic acid leakage the blank microcapsules should turn fluorescent upon completion RT-PCR. We analyzed ∼4.000 randomly picked microcapsules and found no fluorescent events, thus indicating no migration of the target mRNA or cDNA into the microcapsules during RT-PCR assay.

### Accurate differentiation of true positive and false positive events

Accurate differentiation of the fluorescent microcapsules that carry a cell (true positive events) from the fluorescent microcapsules that lack a cell (false positive events) might be challenging since the end-point fluorescent readout can be indistinguishable between these events. We reasoned that using mild-lysis conditions the cell nucleus will retain its compact structure and as a result could be used as a reference to correctly identify the microcapsules with the cells irrespectively of their transcriptional activity. To illustrate this approach, we loaded the mixture of K-562 and HEK293 cells in microcapsules, fixed them in ice-cold ethanol, permeabilized cells in mild-lysis conditions and performed multiplex RT-PCR (see *Materials and Methods*). We confirmed that our approach for freezing and preservation of ethanol-fixed cells did not impact the RT-PCR accuracy, albeit the fluorescence signal intensity was slightly reduced for preserved samples. In addition, we compared the RT-PCR output of ACTB, PTPRC and YAP markers when encapsulated cells were disrupted using mild- and harsh-lysis conditions, and found no significant difference between the two cell-lysis protocols.

Digital analysis of post-RT-PCR samples revealed that using mild-lysis conditions the cell nuclei not only remained compact but also emitted a characteristic, spatially confined fluorescence in all channels (ACTB, YAP, PTPRC) presumably due to non-specific incorporation of fluorescently labelled multiplex PCR probes (**Figure 5B)**. The fluorescence emitting from the nuclei could be enhanced by adding DAPI dye. In contrast, the fluorescence of the PCR amplicons was uniformly distributed across the entire volume of the microcapsule. Therefore, measuring the fluorescence profile of the entire microcapsule provides a straightforward approach to correctly identify the partitions that carry cells and separate them from the partitions with cell-free nucleic acid molecules (**Figure 5C)**. Taking advantage of this feature, we evaluated the cell-free RNA levels of a freshly prepared mixture of cells targeting different house-keeping genes. We anticipated that highly-abundant transcripts such as ACTB would result in increased contamination levels as more RNA molecules will be released prematurely by the compromised cells. In agreement with this notion, we found that the number of cell-free microcapsules displaying a fluorescent signal constituted 16% of positive counts, but dropped down to 0.8% when targeting the TBP gene, which is being expressed at ∼10 copies per cell ^28^. Furthermore, being able to correctly identify and quantify the true/false positives and true/false negatives, we found the multiplex RT-PCR assay specificity to be in the range of 97.71–99.93%. The sensitivity was estimated to be 98.65% for ACTB and 92.57% for YAP on HEK293 cells, and 98.37% for ACTB and 71.74% for PTPRC on K-562 cells. The cell-type specific markers, YAP and PTPRC, displayed excellent positive predictive values (PPVs), 98.56 and 99.25%, respectively. Likewise, negative predictive values (NPVs) were also very high 99.60 and 98.23%, respectively.

**Figure 5.**
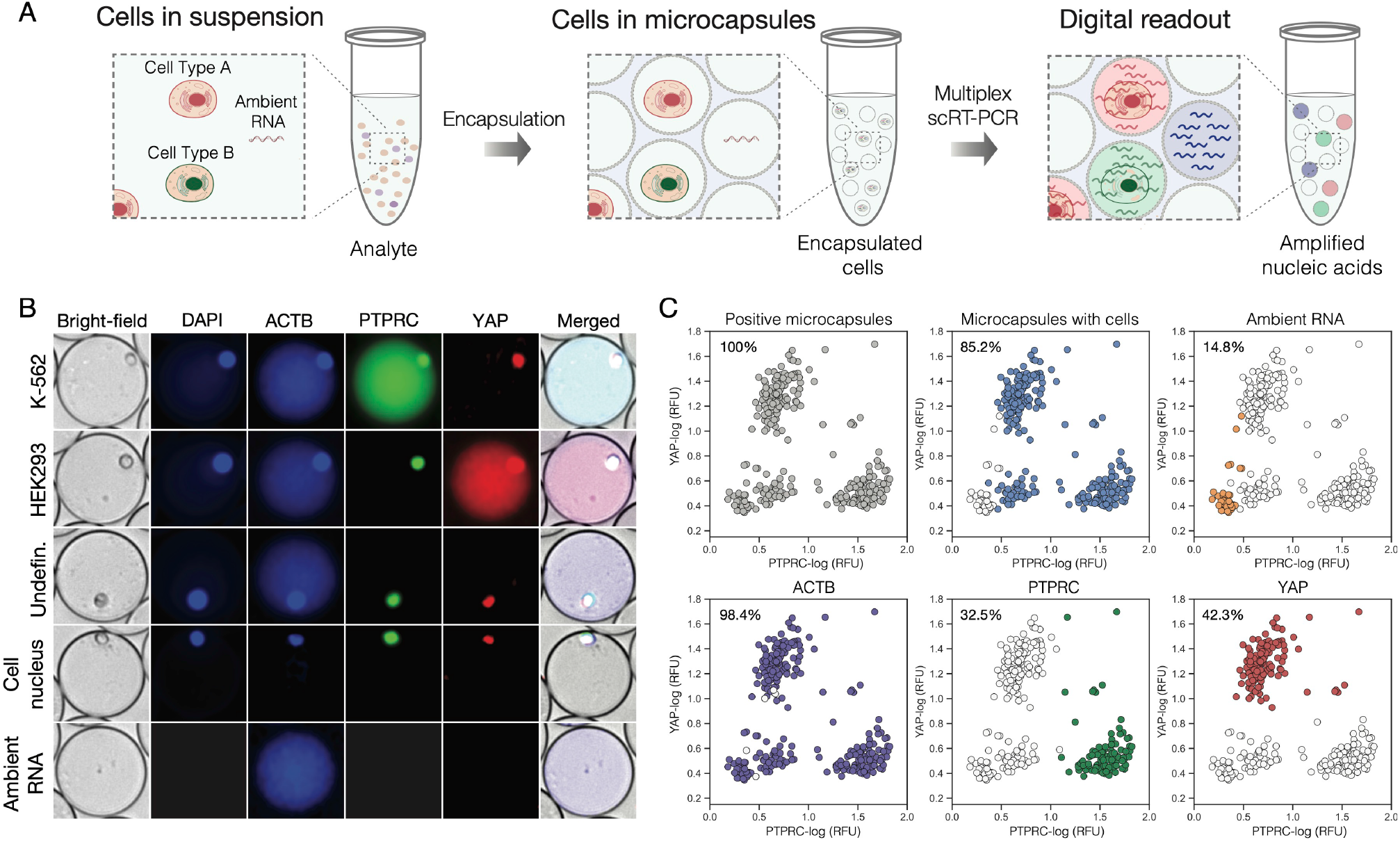
Detection of true positive and false positive events using microcapsule-based multiplex RT-PCR assay. A) Depiction of an experimental scheme where a typical biological analyte comprising cells and ambient RNA is partitioned in microcapsules, subjected to multiplex RT-PCR and evaluated by fluorescence imaging. B) Representative microscopy images of the microcapsules after multiplex scRT-PCR targeting ACTB, PTPRC and YAP markers. Note, the cell nucleus exhibits spatially localized fluorescence in all channels, whereas the PCR amplicons show diffusive fluorescence signal occupying the entire core of a microcapsule. C) Image-based characterization of a biological sample comprising a mixture of K-562 and HEK293 cells. Grey color depicts microcapsules that exhibit a fluorescent signal; blue – microcapsules that contain a cell; orange – microcapsules that exhibit RT-PCR signal lacking a cell; purple, green and red colors depict microcapsules positive for ACTB, PTPRC and YAP signal, respectively. Each plot is represented by >300 counts.

Interestingly, digital profiling of cell mixture uncovered a subpopulation of cells that lacked detectable levels of PTPRC or YAP marker, yet expressed high levels of a house-keeping gene. To better understand the origin of this unusual gene expression signature we performed a separate set of experiments, where multiplex scRT-PCR was performed on each cell type independently. In concordance with FACS ^50^, these experiments revealed a bimodal gene expression distribution in K-562 cells, which are known to differentiate into the erythroid lineage that lacks PTPRC ^51^. However, bimodality may also arise due to transcriptional bursts ^52^, cell cycle dependence ^53^, and stochastic effects ^54^ amongst others, all of which may influence the appearance of cells that lack a transcript of interest at a given moment in time. Therefore, microcapsule-based scRT-PCR output is highly sensitive to the transcriptional state of a cell and captures the individual gene activity at molecular level.

### Identification of leukemia cells in the mixture of peripheral blood mononuclear cells

To further demonstrate the potential use of the developed microcapsule technology tailored for biomedical applications we sought to digitally profile primary human cells. To this end we encapsulated previously-frozen peripheral blood mononuclear cells (PBMCs) and performed scRT-PCR targeting commonly used marker PTPRC (CD45). While we initially expected to achieve >95% detection rate, the image-based as well as FACS-based analysis of the post-RT-PCR microcapsules revealed 75.4-78.6% of the encapsulated cells being positive for the PTPRC. Profiling PBMCs for another ubiquitous marker (B2M) showed similar detection rate. These numbers, however, matched very closely the fraction of viable cells (∼77%), as determined by trypan blue staining, pointing out that cell detection rate by scRT-PCR correlates with cell viability. We then spiked PMBCs with acute promyelocytic leukemia cells (NB-4) and attempted to identify the leukemic cells based on the expression of the fused gene PML-RARα. As it was shown previously, weak PML-RARα expression in the NB-4 cells is within the range detected in primary leukemia samples ^55^. We applied single-cell multiplex RT-PCR as an analogy to the standard clinical practice where instead of individual cells, the total RNA is being extracted from the blood and subjected to cancer diagnostics by bulk RT-PCR ^55,56^. The results presented in **Figure 6** indicate that individual leukemic cells were successfully identified in the mixture of PBMCs, albeit the fraction of detected NB-4 cells appears ∼2-fold lower than the theoretical projections. To better understand this deviation, we quantified the expression levels of PML-RARα transcript in NB-4 cells using qPCR and found it to be ∼200-fold lower than for PTPRC or TBP, which roughly translates to less than 1 copy of PML-RARα transcript per cell, on average. Therefore, it is reasonable to assume that not all NB-4 cells are expressing PML-RARα fusion at a given moment in time, thus explaining the difference in the observed counts. In this context, it is worth noting that digital profiling of another fusion transcript (BCR-ABL) in K-562, which is expressed at approximately 40 copies per cell on average ^57^, resulted in 98.6% detection rate, and targeting transcripts expressed at ∼10 copies per cell (e.g., TBP ^28^) resulted in 97.54% detection rate. Altogether, the RNA cytometry concept based on the semi-permeable microcapsules presented in this work offers a simple, easily customizable approach for rapid and highly sensitive digital profiling of thousands of individual cells at ultra-high-throughput rates, and is advantageous over existing microfluidic and plate-based RT-PCR platforms.

**Figure 6.**
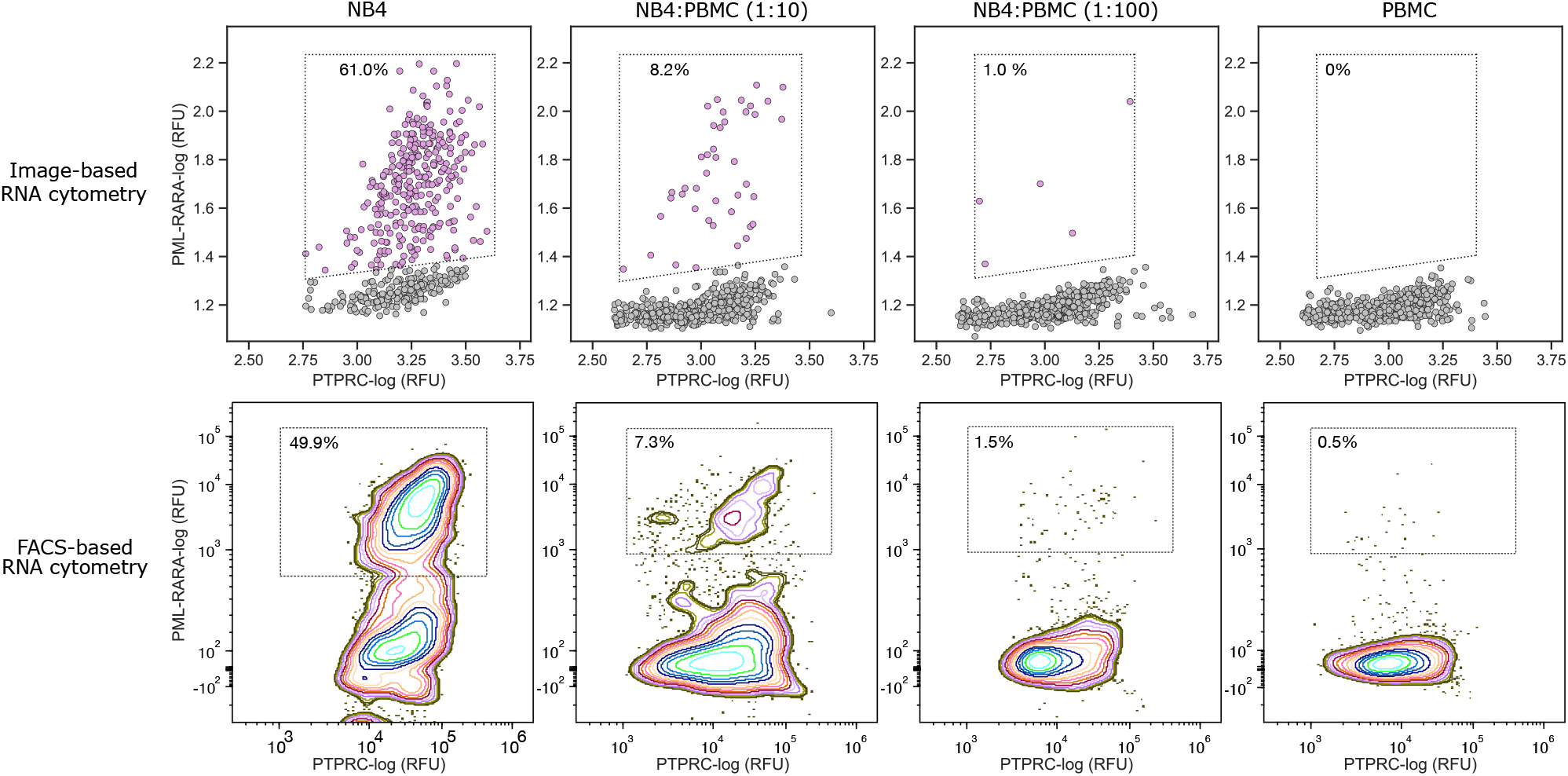
RNA cytometry of acute promyelocytic leukemia cells and human peripheral blood mononuclear cells. A) Image-based and B) FACS-based cytometry of human PBMC spiked with different dilutions of NB-4 cells. The multiplex scRT-PCR targeted PTPRC (CD45) and PML-RARα transcripts. The PBMC cells were detected by expression of marker gene PTPRC, the NB-4 cells were quantified by recording the co-expression of PTPRC and PML-RARα. The percentages depict the fraction of total counts. Refer to accompanying Supplementary Material for gating strategy.

## DISCUSSION

Over the past few years, single-cell gene expression analysis has become one of the essential tools for studying complex biological systems. Single-cell RNA-Seq technology has been particularly instrumental in delineating the cellular functions and discovery of novel biomarkers ^37,58-60^. However, the scRNA-Seq approach still remains a relatively slow and expensive process, and runs counter to the situations built on a fast and accurate assessment of well-characterized genetic markers. For example, assessing the expression of clinically relevant biomarkers using flow cytometry and RT-PCR remains a standard procedure for blood cancers ^55,56,61,62^ and can be completed within a day. While flow cytometry can efficiently discriminate individual cells based on their surface markers, the vast majority of RT-PCR assays, however, employ bulk analyses, which blur the genotypic differences between the cells, and masks clinically relevant cell populations. Therefore, a fast and accurate digital assay for profiling single-cells based on their gene activity provides a valuable and effective option for characterizing heterogeneous biological samples in biomedical research.

In this work we describe, for the first time, a high-throughput RNA cytometry in semi-permeable microcapsules (shells) that enables digital gene expression profiling of tens of thousands of single-cells in a matter of 4 to 6 hours. The biocompatible microcapsules, comprised of a liquid core with dextran and a thin membrane made of covalently cross-linked oligopeptide, efficiently retain the encapsulated cells and nucleic acids during multi-step procedures. The microcapsules remain intact under different conditions, when frozen, thawed or thermocycled, and in the presence of various salts and solvents (e.g., 4M guanidinium isothiocyanate, 90% methanol, 90% acetone). The semi-permeable membrane supports the passive exchange of assay reagents such as enzymes and oligonucleotides, yet simultaneously prevents larger biomolecules such as mRNA, cDNA or gDNA molecules from escaping the microcapsules. We showed that the microcapsules efficiently retain dsDNA molecules longer than 300 bp. at broad range of different temperatures, yet remain permeable to shorter DNA fragments (Figure 2). Strikingly, the post-RT-PCR microcapsules stored at 4 ºC for 10 months retained PCR amplicons, as confirmed by epifluorescence microscopy. While the current permeability cut-off is well suited for conducting a large variety of molecular biology assays, the permeability of the microcapsule’s membrane may be further tuned by altering the cross-linking conditions, adjusting the concentration and modification degree of oligopeptides, thereby providing flexibility for creating microcapsules tailored for specific biological and biomedical applications. This selective permeability simplifies the implementation and use of sequential biochemical reactions on encapsulated species (e.g., cells, microorganisms, viruses, nucleic acids) since the desirable enzymes and reagents can be loaded, or replaced, in the microcapsule by simply changing the reaction buffer that holds the microcapsules. Finally, the complete release of encapsulated nucleic acids is achieved by simply exposing the microcapsules to protease enzyme, which hydrolyzes the peptide bonds.

We demonstrate the utility of microcapsule-based RNA cytometry approach via digital gene expression profiling of human leukemia cells, embryonic kidney cells and primary PBMC using multiplex scRT-PCR assay. At first, we applied harsh-lysis conditions to obtain the microcapsules with a purified total RNA derived from single cells. Subjecting such microcapsules to multiplex RT-PCR generated bright fluorescent partitions specific to a given cell type. However, since the microcapsules with cells and microcapsules with ambient RNA exhibited nearly the same fluorescence intensity (Figure 3B), obtained results also highlighted the danger in quantifying the cells based on the expression of a single gene marker alone. This limitation could be overcome by targeting two genes simultaneously (e.g., one ubiquitously and one differentially expressed gene) to correctly identify the partitions that carry cells (Figure 3C), yet at the cost of increased assay multiplexing. We then showed that scRT-PCR assay using mild-lysis conditions provides a convenient approach for quantifying the cells based on a characteristic, spatially confined fluorescence emanating from a cell nucleus (Figure 5). As a result, the microcapsules with cells and microcapsules with ambient RNA could be clearly differentiated even when the RT-PCR assay is designed to target a single marker gene.

While the scRT-PCR assay specificity was extremely high (99.12 ± 0.89%) with all target probes used in this work, the sensitivity varied from 71 to 98% depending on the gene marker, cell type and primers used. Further optimization of primer pair sequences, RT-PCR conditions and careful vetting of the marker genes, might improve the assay sensitivity. We found that targeting genes expressed at low to medium levels (10 to 40 copies per cell) reduced the false positive (ambient RNA) levels down to ∼0.8%, while retaining high assay sensitivity (98.09 ± 0.77%). The detection of transcripts that are expressed at very low levels (e.g., 1 copy per cell), may need further optimizations one of which may involve using targeted RT primers ^63^. Nonetheless, the presented results show that multiplex RT-PCR assay based on semi-permeable microcapsules provides a reliable approach to profile single-cells rapidly, with high specificity and sensitivity.

The entire process from the initial cell suspension to digital quantification takes a few hours to complete making microcapsule-based RNA cytometry particularly attractive for rapid testing and diagnostics that require fast turnaround and are based on genotypic/transcriptional signatures of individual cells rather than their surface markers. For example, circulating tumor cells (CTCs) often lack proper surface markers, yet their gene expression signatures have been shown to adequately predict the therapy response in multiple cancers ^64-66^. This also applies to biomarkers for which commercial antibodies are limited (e.g., G protein-coupled receptors), or targets for which antibodies cannot be made (e.g., viral transcripts, lncRNA). Another advantage of the microcapsule-based technique reported here is that nucleic acids encoded by single-cells can be efficiently purified under a variety of lysis conditions leading to improved enzymatic reactions. The ability to entirely replace the reagents and easily alter the content of the microcapsules at any given step in a protocol should make microcapsules broadly applicable to different formats of multiplexing ^67^ including *in situ* hybridization and imaging ^68^.

In this work, the post-RT-PCR microcapsules were analyzed using flow cytometry and epifluorescence microscopy. Both approaches generated similar results. In practice, using a multi-color FACS provides the most appealing option for a high-throughput profiling of the individual cells, yet for samples comprising fewer cells (n < 10.000), the image-based analysis may represent a faster and simpler option. By conducting RNA cytometry of individual cells, we accurately identified different cell types and quantified their distribution in the heterogeneous cell population. When using established state-of-art droplet microfluidics platforms, the so-called “signal rain” artifacts create a significant challenge for correctly setting the detection thresholds for separating true negative from false positive events ^67^. Our flow cytometry results, corroborated by microscopy analyses, show that this problem can be overcome by using a microcapsule-based multiplex fluorescence readout. A multiplex scRT-PCR assay based on three simultaneous readouts, i) expression of a ubiquitous marker, ii) expression of a cell-type specific marker(s), and iii) a cell-based marker that does not rely on RNA (e.g., cell nucleus), not only enables accurate identification of distinct cell types present in a heterogeneous mix, but also provides an absolute cell count estimate irrespectively of the transcriptional state of a cell, and the ability to detect ambient nucleic acid molecules (false positives) present in the analyte.

We believe that the semi-permeable microcapsule concept we developed holds a great potential for a broad range of clinical and biological applications. The multiplex scRT-PCR assay can be designed to target gene isoforms ^69^, differentially expressed markers of clinical relevance ^70^, or genetic aberrations ^55,56,71,72^. It can quantitively assess the distribution and frequency of somatic mutations in a panel of cancer-driving genes and benefit cancer diagnostics ^73-75^. Microcapsule-based nested-PCR could be applied for high-throughput screening of B- and T-cells, and for the next-gen sequencing of the immunoglobulin encoded genes ^18,20,24,26,76,77^. Indeed, the applicability of microcapsules can be extended beyond the RT-PCR measurements. The gDNA or cDNA of single-cells in microcapsules could be barcoded and sequenced to obtain the genome-encoded or transcriptome-encoded information. This could contribute greatly to complex disease characterization where genetic aberrations manifest on both DNA and RNA levels. Moreover, the microcapsules presented in this work may provide a very appealing option for single-cell combinatorial indexing using a split-pool synthesis ^78,79^. Due to a selective permeability, the delivery and removal of short barcoding-DNA oligonucleotides during the sequential nucleic acid barcoding reactions becomes straightforward, while simultaneously ensuring efficient retention of longer (barcoded) nucleic acids inside the microcapsules. In addition, because in microcapsules the genetic material released from the lysed cells can be purified and retained through multiple rounds of treatment and washing, it should provide an advantage over droplet-based and plate-based scRNA-Seq approaches, where cell lysis and mRNA barcoding are typically performed in the same reaction mix under suboptimal conditions, and in the presence of intracellular inhibitors (e.g. RNases). Finally, the possibility to completely replace reaction buffers and exchange reagents at any step in the workflow, should provide further flexibility for performing multi-step molecular barcoding reactions. In conclusion, the semi-permeable microcapsule concept that we developed and report here provides a foundation for a variety of single-cell assays that could be built and innovated upon for various goals. We anticipate that this approach will eventually benefit diagnostics, cell biology and biomedicine, where the rapid turnaround is of foremost importance.

## MATERIALS AND METHODS

### Fabrication and use of microfluidic devices

The polydimethylsiloxane (PDMS) microfluidic devices having microchannels 30 and 40 μm heights were obtained from Droplet Genomics.

### Microcapsule generation and cross-linking

The shell-forming solution comprising 3% (w/v) gelatin methacrylate, GMA dissolved in 1x DPBS was loaded onto a microfluidic device along with core-forming solution comprising 15% (w/v) dextran (MW 500k) dissolved in 1x DPBS was loaded onto a microfluidic device using a 1ml syringe connected via 0.56 mm inner diameter PTFE tubing. The resulting aqueous two-phase system (ATPS) droplets quickly formed a core-shell structure comprising a liquid core enriched in dextran, and a liquid shell enriched in GMA. The ATPS droplets were subjected to a two-step polymerization procedure. At first, the water-in-oil droplets were incubated at 4 °C for 15-30 min to solidify the GMA phase. The resulting intermediate-microcapsules, having a liquid core and physically cross-linked GMA shell, were recovered from the oil phase by applying an emulsion breaker (Droplet Genomics), and released into Cell Washing Buffer (1X DPBS and 0.1% w/v Pluronic F-68). The suspension of intermediate-microcapsules was incubated at room temperature for 5 minutes, mixed with a photo-initiator (0.1% (w/v) lithium phenyl-2,4,6-trimethylbenzoylphosphinate (Sigma-Aldrich) and exposed to low-energy 405 nm light emitting diode (LED) device (Droplet Genomics) for 20 seconds. The resulting microcapsules contained a liquid core enriched in dextran and a thin (3.0 ± 0.2 μm) membrane comprising chemically cross-linked polypeptides. Continuing procedures on ice, photopolymerized microcapsules were rinsed twice in Cell Washing Buffer and subjected to cell lysis or cell fixation in ethanol (see below).

### Preparation of cells

K-562, HEK293, and NB-4 cells were cultured in Iscove’s Modified Dulbecco’s Medium (Gibco), Dulbecco’s Modified Eagle’s Medium (Gibco), and Roswell Park Memorial Institute (RPMI) 1640 Medium (Gibco), respectively, supplemented with 10% fetal bovine serum (Gibco) and 1x penicillin-streptomycin (Gibco) at 37 °C in the presence of 5% CO_2_. Cells were collected from a culture dish, washed two times in ice-cold Cell Washing Buffer (1X DPBS and 0.1% w/v Pluronic F-68), and re-suspended in 15% (w/v) dextran solution at a concentration of 0.1-0.2M cells / 100 μl when 40 μm height microfluidic device was used. All centrifugation steps were performed at 300g for 5 min at 4 °C.

### Preparation of PBMCs

PBMCs (ATCC, PCS-800-011) were thawed from liquid nitrogen using RPMI 1640 Medium supplemented with 10% fetal bovine serum. Thawed cells were washed two times in an ice-cold Cell Washing Buffer and resuspended in 1X DPBS at a concentration of 12M cells/ml. All centrifugation steps were performed at 300g for 7-10 min at 4 °C.

### Cell encapsulation

Cell isolation in microcapsules was performed on a microfluidics platform Onyx (Droplet Genomics) using a microfluidic device having a nozzle 40 μm deep and 40 μm wide. Alternatively, cell encapsulation in microcapsules can be performed on custom-build microfluidics platform such as reported previously ^80^. Typical flow rates used were 250 μl/h for GMA solution, 100 μl/h for dextran solution with cells, and 700 μl/h for droplet stabilization oil (Droplet Genomics). NB-4 and PBMC co-isolation in microcapsules was performed using a microfluidic device having a nozzle 30 μm deep and 20 μm wide. Typical flow rates used were 125 μl/h for GMA solution, 50 μl/h for dextran solution with cells, and 700-800 μl/h for droplet stabilization oil. The dilution of cells was chosen such that majority of microcapsules would contain either 0 or 1 cell (occupancy ∼ 0.1). The encapsulations were performed at room temperature for up to 30 minutes. The encapsulated cells were collected in a 1.5 ml tube prefilled with 200 μl of light mineral oil (Sigma-Aldrich).

### Cell fixation in ethanol

To fix the encapsulated cells, the microcapsules were suspended in 70% (v/v) ice-cold ethanol and stored at -20 °C until further analysis. To rehydrate the fixed cells, the tube with microcapsules was equilibrated on ice for 5 minutes, centrifuged at 2000g for 2 minutes at 4 °C, and then washed once in ice-cold 3X SSC buffer supplemented with 0.04% BSA, 1 mM DTT and 0.2U/μl RiboLock RNase Inhibitor. Cells were permeabilized in a mild-lysis buffer as described below.

### Cell lysis

The harsh-lysis of encapsulated cells was performed by suspending microcapsules in 1 ml GeneJET RNA Purification Kit Lysis Buffer (TFS) supplemented with 40 mM DTT. Microcapsules were washed in GeneJET Lysis Buffer 3-times, with 1 to 5 minutes of incubation between the washes. After lysis, the microcapsules were rinsed 5-times in a Washing Buffer (10 mM Tris-HCl [pH 7.5] with 0.1% (v/v) Triton X-100). During washes, the centrifugation steps were performed at 2000g for 2 minutes at 4 °C.

The mild-lysis of encapsulated cells was performed by suspending microcapsules in 1 ml buffer comprising 10 mM Tris-HCl [pH 7.5], 0.6% (v/v) IGEPAL CA-630, 40 mM DTT and 10 mM EDTA. The microcapsules were incubated at room temperature for 15 minutes, rinsed 3-times in a Washing Buffer, and then added to the RT reaction mix. During washes, the centrifugation steps were performed at 2000g for 2 minutes at 4 °C.

### Genomic DNA depletion

Genomic DNA depletion was performed in a 200 μl DNase I reaction mix containing 100 μl close-packed microcapsule suspension, 10U DNase I (TFS), 40U RiboLock RNase Inhibitor, and 1x DNase I Buffer with MgCl_2_, at 37 °C for 20 minutes. Then, an additional 5U of DNase I enzyme were added and incubated for 10 minutes at 37 °C. The microcapsules were rinsed 3-5-times in Washing Buffer and then subjected to a reverse transcription reaction.

### Reverse transcription

cDNA synthesis was performed in 200 μl Maxima H Minus RT reaction mix comprising 100 μl close-packed microcapsule suspension, 5 μM Oligo(dT)21 Primer (IDT), 0.5 mM dNTP Mix, 1000U Maxima H Minus Reverse Transcriptase, 40U RiboLock RNase Inhibitor and 1x RT Buffer, at 50 °C for 60 minutes. Every 20 minutes microcapsules were briefly dispersed. After cDNA synthesis, the RT enzyme was heat inactivated at 85 °C for 5 minutes. Then, microcapsules were rinsed three times in Washing Buffer and subjected to multiplex PCR. For capturing low abundance transcripts (e.g., PML-RARα), 2.5 μM Oligo(dT)_21_ primer was combined with 2.5 μM random hexamer primer (TFS). The cDNA reaction was initiated by preincubating reaction mixture at room temperature for 10 minutes followed by 50 °C for 60 minutes.

### Multiplex PCR

PCR was performed in a 100 μl reaction mix comprising ∼50 μl close-packed microcapsule suspension, 0.5 μM of each PCR primer (see primer sequences in Supplementary Material), and 1x Phire Tissue Direct PCR Master Mix (TFS). Samples were thermally cycled through the following program: 98 °C (5 min), 98 °C (5 s)/64 °C (5 s)/72 °C (20 s) for 30 cycles, 72 °C (1 min). Following thermal cycling, microcapsules were treated with 100 U Exonuclease I (NEB) for 15 minutes at 37 °C, rinsed three times in Washing Buffer, and used for the subsequent microscopy and FACS analysis. For capturing low abundance transcripts, at first the preamplification was performed by 10-cycles of PCR, then microcapsules were washed twice in Washing Buffer and subjected to 30-cycles PCR using identical conditions as indicated above. The post-PCR microcapsules were treated with 100 U Exonuclease I (NEB) for 15 minutes at 37 °C, rinsed three times in Washing Buffer, and used for the subsequent microscopy and FACS analysis.

### Post-PCR microcapsule staining with DAPI

To identify microcapsules with isolated cells, post-RT-PCR microcapsules were immersed in a Washing Buffer containing 300 nM of 4’,6-Diamidino-2-Phenylindole, Dihydrochloride-DAPI (Invitrogen) and incubated in the dark on ice for 10 minutes. Then microcapsules were washed three times in Washing Buffer and used for the subsequent epifluorescence microscopy and FACS analysis.

### Fluorescence microscopy analysis

The fluorescence intensity of microcapsules was recorded by layering the microcapsules on a standard hemocytometer (Sigma-Aldrich) and imaging under Nikon Eclipse Ti-E microscope with DAPI, GFP, RFP, and Cy5 fluorescence filter sets. Imaging settings were kept the same for each experiment with an exposure time of 400 ms, and the gain value set at 1.0. The microscope objective used for imaging was CFI Plan Fluor 10X (N.A. 0.30, W.D. 16.0 mm). For each analysis at least ten brightfield and fluorescence images (∼200 microcapsules per image), were recorded with Nikon DS-Qi2 digital camera.

### Flow cytometry

The microcapsules were washed twice in Washing Buffer, filtered through a 100 μm size cell strainer (Corning), and loaded onto the Partec CyFlow Space (Figure 4) and BD FACSAria III FACS (Figure 6) instruments. The microcapsules were detected using forward scatter, side scatter, and fluorescence channels. Isolated cells in microcapsules were detected by measuring signal height in the DAPI channel on the BD FACSAria III FACS instrument. Note, that due to the spillover of Alexa Fluor 488 to the Alexa Fluor 555 channel and the imperfect compensation process, the PTPRC-positive population showed increased intensity in Alexa Fluor 555 channel when measurements were performed using Partec CyFlow Space (Figure 4).

### RT-qPCR

To evaluate marker expression in cell lines bulk RT-qPCR was conducted using QuantStudio-1 real-time PCR system (TFS). The total RNA from K-562, HEK293, NB-4, and PBMC were extracted using the GeneJET RNA Purification kit (TFS). The gDNA traces were depleted by RapidOut DNA Removal Kit (TFS). cDNA synthesis was performed in 50 μl Maxima H Minus RT reaction mix comprising 2 μg total RNA, 5 μM Oligo(dT)_21_ primer (IDT), 0.5 mM dNTP Mix (TFS), 250 U Maxima H Minus Reverse Transcriptase (TFS), 100 U RiboLock RNase Inhibitor and 1x RT Buffer (TFS), at 50 °C for 30 minutes. After cDNA synthesis, the RT enzyme was heat inactivated at 85 °C for 5 minutes. Then, cDNA material was diluted 10-fold in nuclease-free water and used directly for qPCR. qPCR was performed in 10 μl reaction volume comprising 2 μl cDNA, 5 μl 2x Maxima SYBR Green/ROX qPCR Master Mix (TFS), and 3 μl of 1 μM forward/reverse primer mix (Supplementary Table S1). Samples were thermally cycled through the following program: 95 °C (10 min), 95 °C (15 s)/60 °C (30 s)/72 °C (60 s) for 40 cycles and the number of threshold cycles (Ct) for each marker gene was recorded (Supplementary Table S2) using software provided with the QuantStudio-1 instrument.

### Double stranded DNA retention in microcapsules

GeneRuler Low Range DNA Ladder (TFS) was mixed with 30% (w/w) dextran at ratio 1:1 and encapsulated using the standard procedure described above. Following DNA encapsulation, the emulsion was transferred to 4 °C for 60 minutes. 30 μl aliquot of an emulsion was removed from the tube, broken and treated with 0.5 μl of 20 mg/ml proteinase K (TFS) for 10 minutes at 37 °C and then 10 μl was combined with 2 μl of Gel Loading Dye, Purple (6x) (NEB) and analyzed on 3% agarose gel in 1x TAE buffer (Supplementary Figure S3). This sample was considered as a control since no DNA loss was expected (Well #1). The remaining (150 μl) of the emulsion was converted to microcapsules as follows. The physically cross-linked (intermediate) microcapsules (Well #2) were released from the water-in-oil emulsion by adding 50 μl of emulsion breaker, washed once in 1 ml ice-cold Cell Washing Buffer, pelleted at 500g for 2 minutes at 4 °C. The 30 μl aliquot of physically cross-linked microcapsules was combined with 1 ml of ice-cold Cell Washing Buffer and incubated on ice for 30 minutes. Then, the microcapsules were rinsed twice in ice-cold Cell Washing Buffer, pelleted by centrifugation, and treated with 0.5 μl of 20 mg/mL proteinase K at 37 °C for 10 minutes. Next, 10 μl of the treated sample was combined with 2 μl of Gel Loading Dye, Purple (6x) and analyzed on 3 % agarose gel in 1x TAE buffer (Well#2). The remaining (physically cross-linked) microcapsules were dispersed in 1 ml of ice-cold Cell Washing Buffer supplemented with photo-initiator (0.1% (w/v) LAP). The intermediate-microcapsules were cross-linked by a 20 second exposure to 405 nm LED device (Droplet Genomics) to obtain the microcapsules having a covalently cross-linked membrane. The microcapsules were rinsed once in ice-cold Cell Washing Buffer and then divided into 4 tubes at equal 30 μl portions. 1 ml of Cell Washing Buffer was added to each tube, and microcapsule suspensions were incubated for 30 minutes at different temperatures: 4 °C (Well #3), 22 °C (Well #4), 50 °C (Well #5), and 70 °C (Well #6). Then, the microcapsules that were incubated on ice (4 °C) were rinsed twice in ice-cold Cell Washing Buffer and pelleted by centrifugation. The microcapsules that were incubated at 22-70 °C were rinsed twice in room temperature Cell Washing Buffer and pelleted by centrifugation. The 30 μl of microcapsule suspension of each tube was treated with 0.5 μl of 20 mg/mL proteinase K and incubated at 37 °C for 10 minutes. Finally, 10 μl of each treated sample was combined with 2 μl of Gel Loading Dye, Purple (6x) and analyzed on 3% agarose gel in 1x TAE buffer.

### RNA leakage among microcapsules

To investigate RNA leakage, blank microcapsules (without cells) were immersed in 200 μl Maxima H Minus RT reaction mixture comprising 5 μM Oligo(dT)_21_ Primer (IDT), 0.5 mM dNTP Mix, 1000 U Maxima H Minus Reverse Transcriptase, 40U RiboLock RNase Inhibitor, 1x RT Buffer, 1 μg total RNA purified from K-562 cells and 1 μg total RNA purified from HEK293 cells. Microcapsules occupied half of the final reaction volume (100 μl). RT step was conducted at 50 °C for 60 minutes. Every 20 minutes, microcapsules were briefly dispersed. After cDNA synthesis, the RT enzyme was heat inactivated at 85 °C for 5 minutes. Then, microcapsules were rinsed three times in Washing Buffer and subjected to multiplex PCR targeting ACTB, PTPRC, and YAP. The post-RT-PCR microcapsules (n ∼ 4000) were analyzed under the epifluorescence microscope.

## Data analysis

### Cell retention in microcapsules

To evaluate K-562 cell encapsulation and retention, 36 digital images were recorded: 18 images for water-in-oil droplets and 18 images for microcapsules. Each image contained at least 100 compartments. For each image, the occupancy (lambda value, λ) of cells was estimated. Then, distribution normality was verified by applying Lilliefors (Kolmogorov-Smirnov) normality test. The calculated p-value was 0.6481, which confirmed a normal data distribution. Then F-test for homogeneity of variance was applied. The calculated p-value was 0.372. Assuming equal variance between cell occupancy in droplets and microcapsules, an independent sample t-test was performed (under a two-sided alternative hypothesis). Occupancy measurements were visualized using Python 3.7.6, Pandas framework, and Seaborn library.

### DNA retention in microcapsules

DNA fragment retention was quantified by measuring the DNA band intensity on an agarose gel. Each DNA fragment retention value was averaged from three independent measurements and analyzed using a Fiji software package. The measurements were visualized using Python 3.7.6, Pandas framework, and Seaborn library.

### Microscopy analysis of post-RT-PCR microcapsules

Microcapsules from bright-field images were analyzed using a Python script (provided in the GitHub repository). Microcapsules were outlined using Hough circle transform and the masks were used to crop and measure mean and max fluorescence from corresponding DAPI, GFP, RFP, and Cy5 filtered images. Measurements were processed and visualized using Python 3.7.6, Pandas framework, and Seaborn library.

### Flow cytometry data analysis

Flow cytometry data were analyzed and visualized using FCS Express 7 software (version 7.12.0005). The post-RT-PCR microcapsules after harsh-lysis were analyzed using the Partec CyFlow Space FACS instruments. The gating process was performed in the following manner: 1) gating the microcapsules based on forward vs. side scatter signal, 2) gating ACTB positive events based on Alexa Fluor 647 vs. side scatter signal, 3) analyzing PTPRC and YAP marker abundance based on Alexa Fluor 488 vs. Alexa Fluor 555 signal. All measurements were performed by analyzing the signal area.

The post-RT-PCR microcapsules after mild-lysis on NB-4 and PBMCs were analyzed using the BD FACSAria III FACS instrument. The gating process was performed in the following manner: 1) gating the microcapsules with isolated cells based on Alexa Fluor 488 signal area vs. DAPI-stained nuclei fluorescence signal height, 2) gating PML-RARα positive events based on Alexa Fluor 488 signal area vs. Alexa Fluor 555 signal area.

### Sensitivity, specificity, and positive/negative predictive values

Sensitivity was defined as *Sensitivity* = (*TruePos*/(*TruePos* + *FalseNeg*)) × 100%. Specificity was defined as *Specificity* = (*TureNeg*/(*TureNeg* + *FalsePos*)) × 100%. Positive predictive value was defined as *PPV* = (*TruePos*/(*TruePos* + *FalsePos*)) × 100%. Negative predictive value was defined as *NPV* = (*TureNeg*/(*TureNeg* + *FalseNeg*)) × 100%. The microcapsules carrying a cell and being positive for one of the marker genes (ACTB, B2M, TBP, PTPRC, YAP) were counted as true positives. The microcapsules carrying no cells and showing no fluorescence were counted as true negatives. The false positives were microcapsules lacking cells but being fluorescent for one of the marker genes. The false negatives were microcapsules with cells displaying no signal for any marker gene.

## ACKNOWLEDGEMENTS

This work has received funding from European Regional Development Fund (project No 01.2.2-LMT-K-718-04-0002]) under grant agreement with the Research Council of Lithuania. Authors are especially grateful to Dr. Karolis Leonavičius (Droplet Genomics, Lithuania) for image analysis consulting and assistance, Indrė Dalgėdienė (Vilnius University, Lithuania) for assistance with Partec CyFlow Space FACS instrument and Dr. Vytautas Kašėta (CIM, Lithuania) for assistance with BD FACSAria III FACS instrument. Authors are grateful to Jonas Gasparavičius (Droplet Genomics, Lithuania) for supplying microfluidic devices used in this work, Suji Kim (Vilnius University, Lithuania) for assistance with cell preparation and Dr. Veronika Viktorija Borutinskaitė (Vilnius University, Lithuania) for kindly providing NB-4 cell line. Authors express their gratitude to Arūnas Stirkė (FTMC, Lithuania) for kindly providing an access to epifluorescence microscope.

